# Genomic evidence for inter-class host transition between abundant streamlined heterotrophs by a novel and ubiquitous marine Methylophage

**DOI:** 10.1101/2021.08.24.457595

**Authors:** Holger H. Buchholz, Luis M. Bolaños, Ashley G. Bell, Michelle L. Michelsen, Michael J. Allen, Ben Temperton

## Abstract

The methylotrophic OM43 clade are Gammaproteobacteria that comprise some of the smallest free-living cells known and have highly streamlined genomes. OM43 represents an important microbial link 0between marine primary production and remineralisation of carbon back to the atmosphere. Bacteriophages shape microbial communities and are major drivers of microbial mortality and global marine biogeochemistry. Recent cultivation efforts have brought the first viruses infecting members of the OM43 clade into culture. Here we characterize a novel myophage infecting OM43 called Melnitz. Melnitz was isolated independently on three separate occasions (with isolates sharing >99.95% average nucleotide identity) from water samples from a subtropical ocean gyre (Sargasso Sea) and temperate coastal (Western English Channel) systems. Metagenomic recruitment from global ocean viromes confirmed that Melnitz is globally ubiquitous, congruent with patterns of host abundance. Bacteria with streamlined genomes such as OM43 and the globally dominant SAR11 clade use riboswitches as an efficient method to regulate metabolism. Melnitz encodes a two-piece tmRNA (*ssrA*), controlled by a glutamine riboswitch, providing evidence that riboswitch use also occurs for regulation during phage infection of streamlined heterotrophs. Virally encoded tRNAs and *ssrA* found in Melnitz were phylogenetically more closely related to those found within the alphaproteobacterial SAR11 clade and their associated myophages than those within their gammaproteobacterial hosts. This suggests the possibility of an ancestral inter-class host transition event between SAR11 and OM43. Melnitz and a related myophage that infects SAR11 were unable to infect hosts of the SAR11 and OM43, respectively, suggesting host transition rather than a broadening of host range.

**IMPORTANCE:** Isolation and cultivation of viruses is the foundation on which the mechanistic understanding of virus-host interactions and ground-truthing is based. This study isolated and characterised the first myophage known to infect the OM43 clade, expanding our knowledge of this understudied group of microbes. The near-identical genomes of four strains of Melnitz isolated from different marine provinces and global abundance estimations from metagenomic data suggest that this viral population is globally ubiquitous. Genome analysis revealed several unusual features in Melnitz and related genomes recovered from viromes, such as a curli operon and virally encoded tmRNA controlled by a glutamine riboswitch, neither of which are found in the host. Further phylogenetic analysis of shared genes indicates that this group of viruses infecting the gammaproteobacterial OM43 shares a recent common ancestor with viruses infecting the abundant alphaproteobacterial SAR11 clade. Host ranges are affected by compatible cell surface receptors, successful circumvention of superinfection exclusion systems and the presence of required accessory proteins, which typically limits phages to singular narrow groups of closely related bacterial hosts. This study provides intriguing evidence that for streamlined heterotrophic bacteria, virus-host transitioning is not necessarily restricted to phylogenetically related hosts, but is a function of shared physical and biochemical properties of the cell.

## Introduction

Bacteriophages are the most abundant and diverse biological entities in the oceans and are, on average, an order of magnitude more abundant than their bacterial hosts in surface water (1, 2). Viral predation kills a large proportion of bacterial cells in marine surface waters each day (3) and is a main contributor to nutrient recycling by releasing cell-bound organic compounds into the environment (4, 5). Viral infection can also alter host metabolism through metabolic hijacking (6, 7), which has been shown to reprogram resource acquisition and central carbon and energy metabolism (8), influencing oceanic nutrient cycles. The selective pressure of the predator-prey relationship of bacteria and phages is also a main driver of microbial evolution (9), where the constant arms race requires phages to evolve and improve strategies to overcome host defence mechanisms (10). Recent advances in culture-independent sequencing technology such as single-cell genomics and metagenomics have expanded our understanding of the enormous diversity of marine viruses (11–14). However, many of these sequences lack representation in viral culture collections (15), limiting experimental determination of parameters of infection such as host-range. A resurgence in bacterial cultivation efforts and improved viral isolation methods has led to the discovery of many new phages infecting abundant but fastidious marine bacteria such as SAR11. Combining genomes from viral cultures with metagenomics identified these viruses to be some of the most abundant on Earth across all marine ecosystems (15–18). Yet, many more virus-host systems occupying a range of important ecological niches such as methylotrophy remain poorly understood.

Members of the OM43 clade are small, genomically streamlined (genomes ∼1.3 Mbp) Type I methylotrophs of the class Gammaproteobacteria (19). The catabolism of methanol and other volatile organic compounds (VOCs) is an important link between primary production and remineralisation of carbon back to atmospheric CO_2_ (20–22). In the surface ocean, peak abundance of OM43 coincides with phytoplankton blooms which provide their main carbon source (22). OM43 are particularly abundant in coastal ecosystems where they comprise up to 5% of the microbial community (23). Members of the OM43 clade are somewhat challenging to grow in the laboratory: increased levels of auxotrophy and largely constitutive metabolism renders them sensitive to media composition (19). As a result, only two OM43 phages have been reported, therefore the influence of viral predation on OM43 is virtually unexplored. The isolation of the first viruses infecting OM43 (Venkman) from the coastal Western English Channel (WEC); and MEP301 from the Bohai Sea were both reported in 2021 (15)(24). Venkman was the third most abundant phage in the WEC sample, indicating that phages of methylotrophs are a major component of this coastal ecosystem (15). In contrast, recruiting reads from global ocean viromes against MEP301 and Venkman, indicated that their relative abundance was below detection limit in most lower-latitude pelagic viromes (24). Thus, phages infecting OM43 were thought to be predominantly found at higher latitudes in regions of high primary productivity.

Here we report the isolation and genomic analysis of Melnitz, representing a novel population of myophages infecting OM43. Four representatives of this virus that shared >99.5% average nucleotide identity were isolated independently on three separate occasions from the temperate coastal WEC and from the Sargasso Sea located within the North Atlantic subtropical gyre. This indicates that despite low relative abundance at low latitudes, Melnitz was sufficiently abundant to be isolated through enrichment techniques. The genomic similarity between independent isolates suggests a cosmopolitan global distribution, which was supported by metagenomic read recruitment from global ocean viromes. Genome analysis of Melnitz revealed a two-piece tmRNA gene (*ssrA*), controlled by a glutamine riboswitch. Riboswitch control of regulation is a feature of streamlined organisms such as OM43 and SAR11 (25), and here we show that it is also a feature of their associated viruses. Like previously reported SAR11 myophages (15, 16), Melnitz also encoded the portal proteins of a curli operon that is absent in the host. Structural analysis suggests a putative reconfiguration allows this phage-encoded protein to serve as a novel gated secretin or pinholin, with gene synteny indicating a role in timing the release of viral progeny. Phylogenetic analysis revealed that both the *ssrA* gene and tRNA genes encoded by Melnitz were more closely related to those found within the alphaproteobacterial SAR11 host or its associated viruses, than those of its own gammaproteobacterial host. These findings point towards a recent shared ancestor indicative of host transitioning between OM43 and SAR11.

## Results and Discussion

### Phage Melnitz infecting Methylophilales sp. H5P1 shares viral clusters with Pelagibacter phages

Two bacteriophages were isolated on the OM43 strain H5P1 from two environmental water samples taken from the Western English Channel (WEC) previously (15). Two more phages were obtained from an additional water sample taken at the Bermuda Atlantic Time Series (BATS) station in the Sargasso Sea (Table 1). Phages were successfully purified, sequenced from axenic cultures and assembled into single circular contigs. CheckV comparison to publicly available metagenomes and phage genomes suggested that the viral contigs were complete, circularly permuted genomes without terminal repeats. Transmission electron microscopy (TEM) showed straight contractile tails indicative of myophage morphology with a capsid size of 58±6 nm (Supplementary Figure 1).

**Table 1.** General features of OM43 strain H5P1 and four Melnitz phages isolated in this study.

| Phage | Culture ID | Host | Phage group | Morphotype | Genome size (bp) | G + C % | ORFs | tRNAs | Source Water | Sampling Date | Latitude | Longitude |
| --- | --- | --- | --- | --- | --- | --- | --- | --- | --- | --- | --- | --- |
| Melnitz | EXVC044M | H5P1 | Melnitz | Myovirus | 141,548 | 37.6 | 224 | 4 | BATS | 01 June 2019 | N31°40' | W64°10' |
| Melnitz variant 1 | EXVC043M | H5P1 | Melnitz | Myovirus | 141,552 | 37.6 | 226 | 4 | BATS | 01 June 2019 | N31°40' | W64°10' |
| Melnitz variant 2 | EXVC040M | H5P1 | Melnitz | Myovirus | 141,548 | 37.6 | 224 | 4 | WEC | 22 July 2019 | N50°15' | W04°13' |
| Melnitz variant 3 | EXVC039M | H5P1 | Melnitz | Myovirus | 141,372 | 37.6 | 224 | 4 | WEC | 01 April 2019 | N50°15' | W04°13' |
|  |  | H5P1 | Methylophilales sp. | Gammaproteobacteria | 1,336,408 | 34.4 | 1382 | 38 | WEC | Sep-18 | N50°15' | W04°13' |

All four phage genomes shared 99.95-100% average nucleotide identity across their full genomes therefore all four phages should be regarded as the same viral species (26). We named this species “Melnitz’’ after a character in the popular Ghostbusters franchise, continuing the theme of another previously isolated phage on OM43 (Venkman) (15). For clarity, where individual phages within the population (Melnitz) are specified, they will subsequently be referred to with a numerical suffix (e.g. Melnitz-1). General features of the four phages are summarised in Table 1. Shared gene network analysis (VConTACT2) using assembled contigs from Global Ocean Viromes (GOV2) and RefSeq (V88 with ICTV and NCBI taxonomy) viruses assigned Melnitz to the same cluster as the *Pelagibacter* myophages HTVC008M and Mosig (16, 27), suggesting they belong to the same family (Figure 1). The four Melnitz genomes were shared across two clusters, the first contained an additional 15 viral contigs from nine different metagenomes. The second cluster contained three virome contigs shared between the two clusters, as well as 25 pelagimyophage genomes assembled from metagenomes (PMP-MAVGs) (28). *Pelagibacter* myophage isolates HTVC008M and Mosig were also placed in this second cluster. A total of 66 contigs clustering with Melnitz isolates were identified from GOV2 and WEC viromes (12, 29). Phylogenetic analysis showed that only two environmental contigs consistently shared a branch with Melnitz, based on single shared genes encoding: tail sheath protein (Supplementary Figure 2), terminase large subunit (Supplementary Figure 3) and scaffolding proteins (Supplementary Figure 4), as well as four concatenated structural genes (Figure 2). All phages within this viral group were either isolates known to infect streamlined heterotrophs, or were previously predicted to do so based on phylogenetic similarity (28).

**Figure 1.**
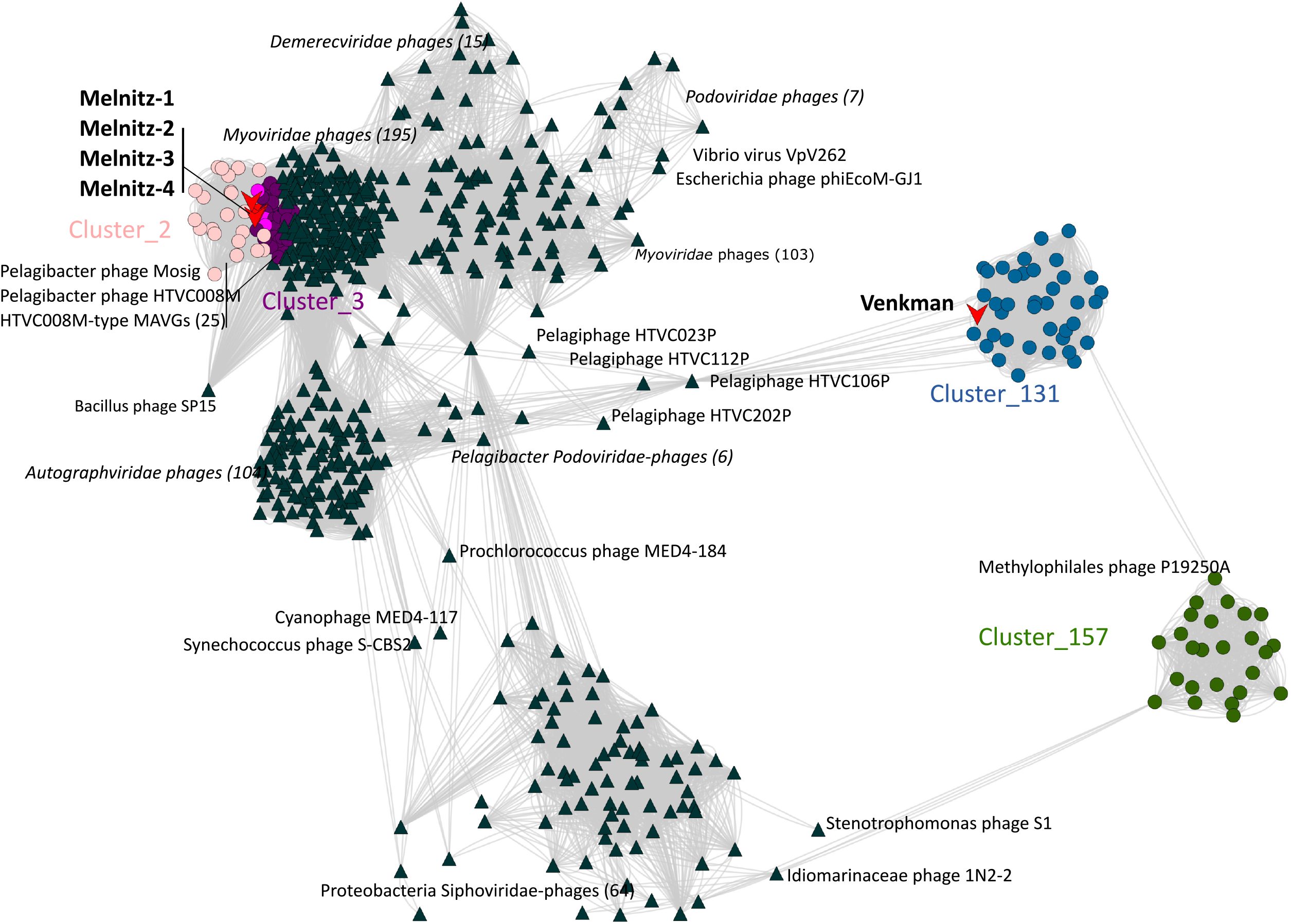
Viral shared gene-content network of OM43 phages, related bacteriophages from NCBI and related sequences from the Global Ocean Virome (GOV2.0). Nodes represent viral genomes; edges represent the similarity between phages based on shared gene content. NCBI reference genomes more than two neighbouring edges removed were excluded for clarity. Phage isolates are indicated with red arrows. Coloured circles represent genomes and virome contigs within the same cluster as OM43 phage isolate genomes. Nodes shared between Cluster_2 (light pink) and Cluster_3 (purple) are highlighted in hot pink.

**Figure 2.**
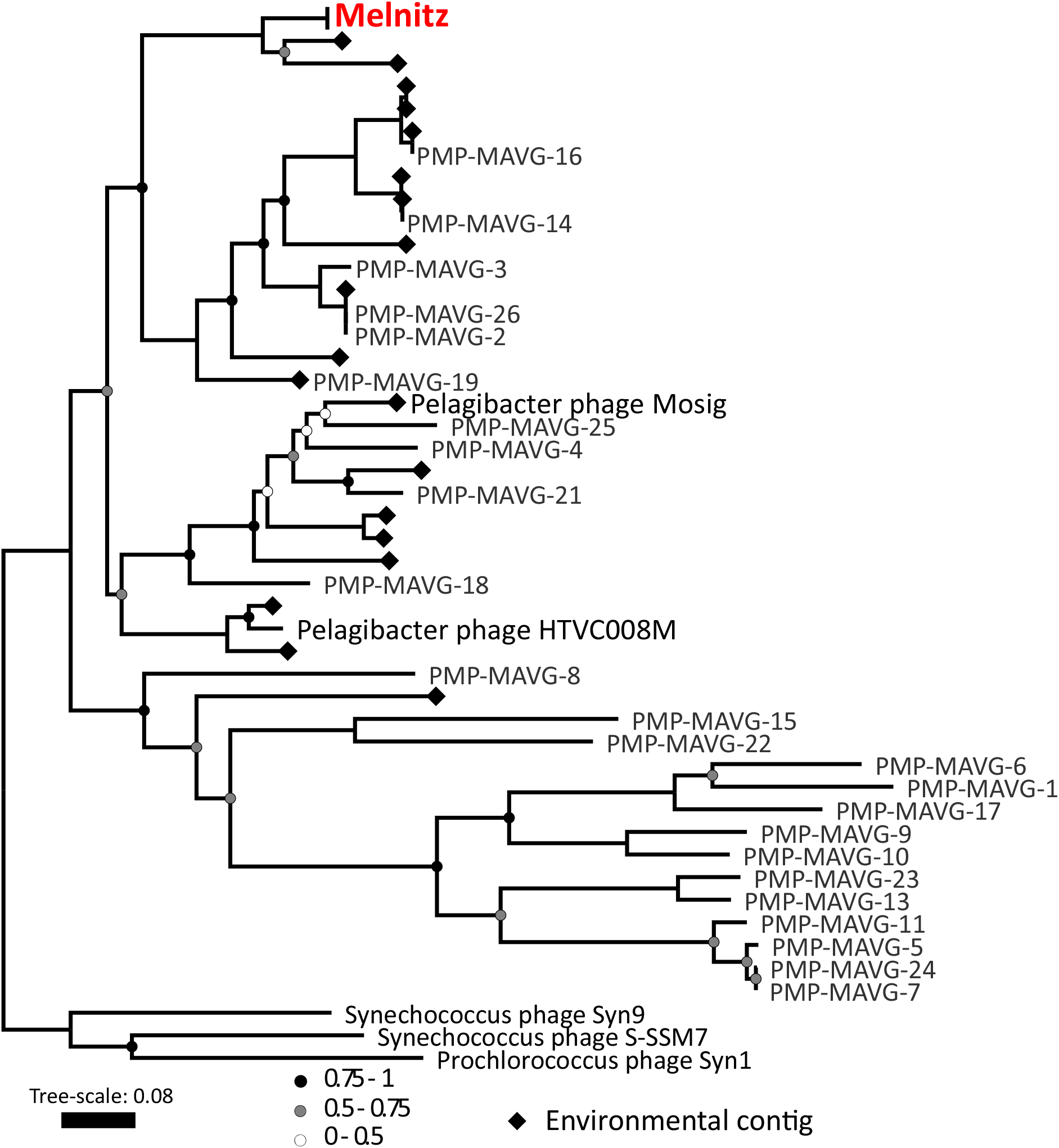
Phylogenetic tree of metagenomic contigs and marine Melnitz-type myophages. Neighbour-joining tree (500 bootstraps) based on four individually aligned and concatenated structural genes (capsid assembly, major capsid, sheath subtilisin, terminase large subunit) of genomes and contigs that were clustered with OM43 phage Melnitz, with the exception of phages infecting *Synechococcus* spp. that were used to root the tree. All four Melnitz-like isolates were included but the branch was collapsed for clarity. Branch support values of 1 are not shown. Leaves without labels indicate contigs from the Global Ocean Virome (GOV2) dataset (12), which were omitted for clarity.

### Metagenomic analysis shows cosmopolitan nature of Melnitz-like phages

To establish the distribution patterns of Melnitz, GOV2 virome reads were mapped against phage genomes (≥95% nucleotide identity over ≥90% read length; genome coverage of <40% was required to register a phage as ‘present’ to avoid false positives (30)) (Supplementary Figure 5). Relative abundance of phages was calculated based on the number of reads mapped to a contig, normalised by contig length and sequencing depth (mapped reads per kilobase pair of metagenome; RPKM). Linear regression of relative abundance of all three phages known to infect OM43 (Melnitz from this study, plus Venkman and MEP301) as a function of temperature showed a significant and negative relationship (Melnitz: *p* = 4.69 × 10^−5^, R^2^ = 0.199; Venkman: *p* < 2.20 × 10^−16^, R^2^ = 0.435; MEP301: *p* = 1.85 × 10^−7^, R^2^ = 0.198). Relative abundance was positively correlated with absolute latitude (Melnitz: p = 1.78 × 10^−5^, R^2^ = 0.094; Venkman: p < 2.20 × 10^−16^, R^2^ = 0.545; MEP301: p = 4.67 × 10^−9^, R^2^ = 0.247). This suggests that these viruses are most abundant in cold, productive waters. In contrast, relative abundances of Melnitz-related *Pelagibacter* phages HTVC008M and Mosig were not correlated to temperature (Mosig: *p*= 0.094, R^2^ = 0.047; HTVC008M: *p*= 0.217, R^2^ = 0.004) or latitude (Mosig: *p*= 0.028, R^2^ = 0.031; HTVC008M: *p*= 0.077, R^2^ = 0.018). OM43 phages MEP301 and Venkman were classed as ‘present’ (>40% genome coverage) in 39.1% and 53.9% of 131 GOV2 viromes, respectively. Melnitz showed greater global ubiquity, being classed as ‘present’ in 78.8% of GOV2 viromes. Whilst there are no GOV2 samples from the Sargasso Sea, this ubiquity may explain how we were able to isolate Melnitz from both the WEC and the Sargasso Sea.

It has been demonstrated that the marine biosphere maintains persistent bacterial “seed banks” (31), meaning there is a high probability that any given marine bacteria can be found in any marine ecosystem, albeit in extremely low abundance, awaiting favourable conditions for growth. Similarly, many viruses infecting globally distributed bacteria such as SAR11 are found in viromes from all oceans (18). Studies using “viral tagging” on cyanobacteria and their associated cyanophages suggest that at single sites, viral populations in metagenomes comprise non-overlapping viral sequence space resembling “clouds”, which has benchmarked metagenomic analyses and subsequent interpretations (32, 33). In that model, single strains within a viral population encode genomic variations that enable the overall population to adapt more rapidly to environmental selection pressures. In the case of Melnitz, the isolation of the same virus with up to 100% nucleotide identity across the full genome on three separate occasions in the Western English Channel and BATS station in the Sargasso Sea (∼5000 km distance between sites) suggests either low population-level variance, or that the isolation conditions used favour this strain. Nonetheless, the presence of cultivable Melnitz populations at both sites supports the virus seed-bank hypothesis where viral populations are conserved and persistent in the environment, being passively transported across oceans via global currents until favourable conditions select them for propagation (34, 35). The ‘environmental selection’ in the case of Melnitz would likely be the enrichment culturing used for isolation, providing enough suitable hosts and nutrients for viral replication. The possibility remains that the Melnitz population at the BATS site is maintained by a resident “seed” population of OM43, though OM43 is seldom reported in microbial communities at BATS.

### Genomic characterization of Melnitz-like Methylophilales phages

Genomes of the four Melnitz-like myophage isolates were between 141,372 - 141,552 bp in length, all with G+C content of 37.6 % (Table 1), similar to that of its host (34.4%). For each, 224 open reading frames (ORFs) were predicted, except Melnitz-2, which had two additional small ORFs of unknown function (226 ORFs total). Four additional tRNA sequences for all four genomes were identified, and functional annotation of ORFs suggest that all Melnitz strains encoded the same set of genes. Out of 224 ORFs, 143 (∼63%) had unknown function (Figure 3A). The tail-assembly associated region in the Melnitz-4 genome had nine genes with altered length compared to the equivalents in the other three phages, but where annotation was possible, the genes were predicted to have the same function. Predicted protein structures (PHYRE2) had low confidence and did not allow for a meaningful structural comparison (data not shown). Though structural variations in tail and receptor genes are often considered to be important factors for defining strain-level host ranges (36, 37), all four Melnitz variants had identical host ranges when screened against a panel of other OM43 isolates from the WEC (Methylophilales sp. C6P1, D12P1 and H5P1) (15) (data not shown). Melnitz possessed a set of structural genes typically associated with *E. coli* T4-type myophages, including T4-like baseplate, tail tubes, base plate wedges, tail fibres, virus neck, tail sheath stabilization, prohead core and capsid proteins. Melnitz encodes orthologs of the auxiliary metabolic genes (AMGs) *mazG* and *phoH*, which are involved in cellular phosphate starvation induced stress responses and are a common feature of phages from P-limited marine environments (38–40). Additionally, *hsp20* was identified, which together with *mazG* and *phoH* are considered part of the core genes in T4-like cyanomyophages (38, 41). Both Melnitz and its host, Methylophilales sp. H5P1, encode for a type II DNA methylase; DNA methylase targets foreign DNA for cleavage by the restriction endonucleases (42). In T-even phages, DAM methylase (similar to type II DNA methylase) protects phage DNA from restriction endonucleases through competitive inhibition (43), thus DNA methylase in Melnitz could also be involved in protection from host restriction endonucleases during infection.

**Figure 3.**
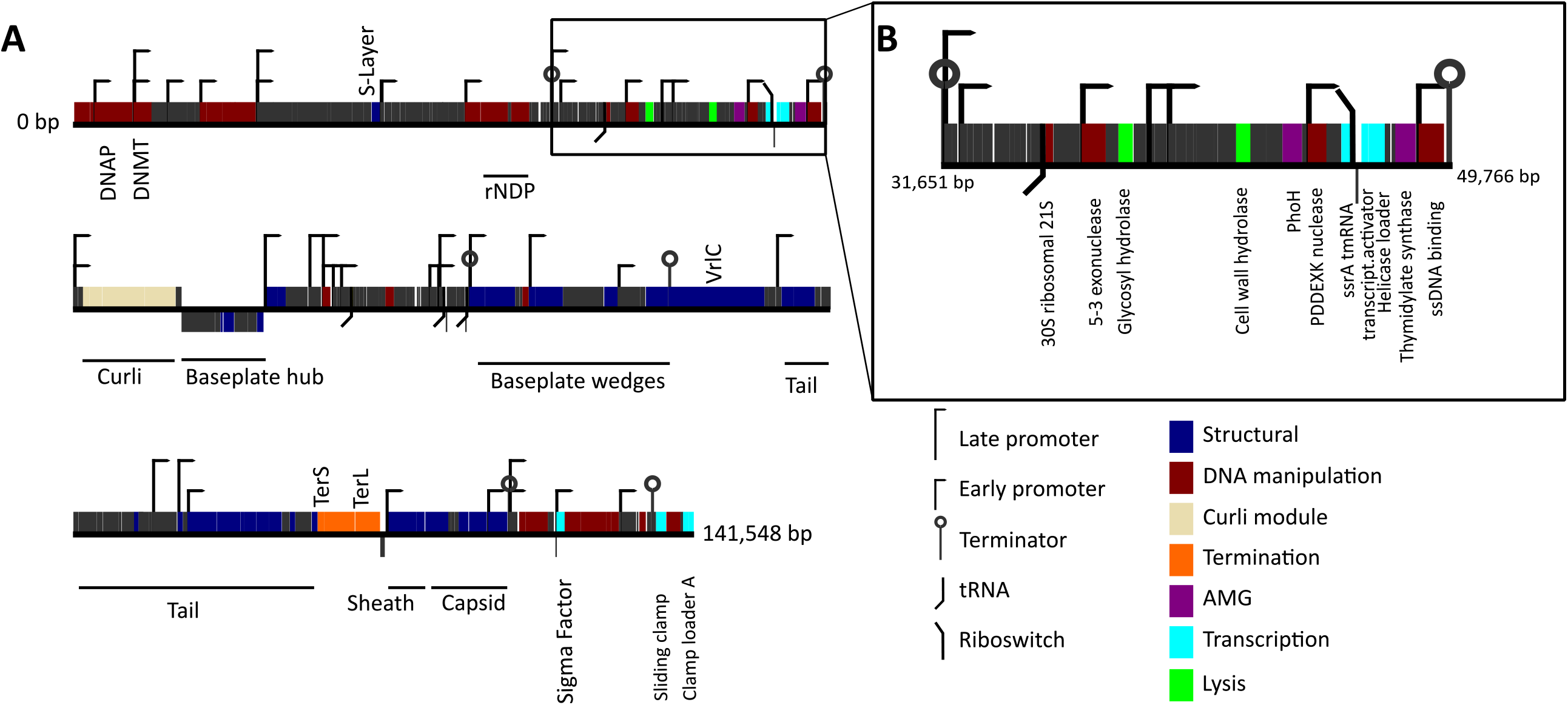
Gene map showing identified genomic features of the OM43 phage Melnitz. **A:** Gene map of the 141,548 bp genome of Melnitz contains 143 hypothetical ORFs (62 %) without known function (indicated dark grey); **B:** Section of the Melnitz genome between two terminator sequences that contains the *ssrA* gene, glutamine riboswitch and transcription co-activator gene for transcription-translation regulation.

For DNA replication and metabolism, Melnitz encodes *nrdA/nrdE* and *nrdB/nrdF* genes that together form active ribonucleotide reductases for catalysing the synthesis of deoxyribonucleoside triphosphates (dNTPs) required for DNA synthesis (44). Additional DNA replication and manipulation associated genes were *polB, recA, dnaB*, as well as two DNA primases and a 2OG-Fe oxygenase. A *rpsU* gene (encoding the bacterial 30S ribosomal subunit 21S) was found downstream adjacent to a tRNA-Arg (tct). Virally encoded S21 genes were previously identified in the *Pelagibacter* phages HTVC008M (16) and Mosig (27) as well as the OM43 phage Venkman (15). It is thought that the 21S subunit in phages might be required for initiating polypeptide synthesis and mediation of mRNA binding (45); the proximate tRNA sequence may support a translational and protein synthesis role of the viral ribosomal gene. Within the same region, Melnitz also encodes for a surface layer-type (S-layer) protein, previously associated with a strong protection mechanism against superinfection in bacteriophages of *Bacillus spp*. and *Pseudomonas aeruginosa* (46–48). We postulate that phage-encoded S-layer proteins in Melnitz may be used to alter host cell surface receptors and thereby protect against superinfection from viral competitors.

Temporal regulation of T4-like phages primarily occurs at the transcriptional level and requires concomitant DNA replication and promoter/terminator sequences to organise into early, middle and late stages (49). In Melnitz, a total of 46 promoters (31 early and 15 late) and six terminator sequences were identified – considerably fewer than the >124 promoter sequences found in T4 (50, 51). Like other marine T4-like phages, such as those infecting streamlined cyanobacteria (52), Melnitz lacked identifiable middle promotors. Reduced numbers of regulatory elements are a common feature of genomically streamlined bacteria such as SAR11 and OM43 (53). Therefore, we propose that temporal regulation of Melnitz is similarly simplified as a result of host streamlining. An alternative hypothesis is that additional (middle-) promoter sequences in Melnitz are too divergent from known sequences for detection.

Like other T4-like phages, Melnitz did not encode for RNA polymerase, relying on host transcription machinery after infection for synthesis of phage proteins. Melnitz possessed the essential T4-like transcriptional genes: σ^70^-like; gp45-like sliding clamp C terminal; clamp loader A subunit; transcription coactivator; and a translational regulator protein. In T4, the DNA sliding clamp subunit of a DNA polymerase holoenzyme coordinates genome replication in late-stage infection, forming an initiation complex with a sigma-factor protein and host RNA polymerase (54). To activate it, the sliding clamp is loaded by a clamp-loader DNA polymerase protein complex, allowing the complex to move along DNA strands (55, 56). As the same genes were found in Melnitz it is likely using them to form a T4-like protein complex for transcription and genome replication.

### Marine Melnitz-like phages may use glutamine riboswitches to regulate genome expression

*ssrA* encodes for a tmRNA that in bacterial *trans*-translation together with *smpB* and ribosomal protein S1 is important to release ribosomes that have stalled during protein biosynthesis. These proteins are subsequently tagged and degraded (57). In Melnitz, the *ssrA* gene encoding two-piece tmRNA was situated on the same operon and directly upstream of the transcription co-activator (Figure 3B). Host-encoded *ssrA* can be used as sites of integration for prophages in deep-sea *Sherwanella* isolates (47) and fragments of *ssrA* have previously been identified in prophage genomes (48), most likely as a result of imprecise prophage excision that transfers host genetic material to the excising phage. Though we did not find integrases or other evidence that Melnitz is able to integrate into host genomes, a speculative temperate ancestral strain may explain the presence of a complete *ssrA* gene in Melnitz. A possible role for a complete viral tmRNA is to either help maintain the hijacked bacterial machinery, or the phage might use tmRNA to selectively tag and degrade host proteins to recycle amino acids for viral protein synthesis. We evaluated the frequency that *ssrA* genes were found in 18,146 publicly available genomes in the phage genome database at millardlab.org (last updated on January 21, 2021), which includes all RefSeq genomes. Only 402 phages (2.3% of all available genomes) encoded 133 unique tmRNA genetic structures. Of these, 53 were suggested to be two-piece tmRNA by ARAGORN (59), of which 50 were encoded by phages isolated from marine or aquatic samples. In contrast, only 2.5% of non-permuted tmRNAs were marine, suggesting that viral permuted tmRNA is a feature more prevalent in marine and aquatic phages compared to phages from other environments. Of the 47 contigs (27 were confirmed complete) that were clustered with Melnitz based on shared gene content (Figure 1), 17 had complete *ssrA* genes, indicating that *ssrA* encoded tmRNA is a common, but not defining feature of this viral group.

The 3’ domain of the *ssrA* gene in Melnitz encodes a predicted glutamine riboswitch that resembles *glnA* RNA motifs found in cyanobacteria and other marine bacteria (58). Riboswitches are a common regulatory mechanism in streamlined marine bacteria due to their low metabolic maintenance cost compared to protein-encoded promotors and repressors (53). Like tmRNAs, riboswitches are a rare feature in phages. A phage-encoded riboswitch putatively controlling regulation of *psbA* was previously identified in a cyanophage (56). Of the 133 isolate phage genomes encoding tmRNAs only two phages other than Melnitz possessed a riboswitch (both glutamine): *Pelagibacter* phage Mosig (27) and *Prochlorococcus* phage AG-345-P14 (60). Additional riboswitches were identified in metagenomic assembled virome contigs. In ten metagenomically assembled viral genomes of pelagimyophages thought to infect SAR11 (PMP-MAVGs) and five GOV2 contigs related to Melnitz, only one of these contigs had both tmRNA and a riboswitch; eleven PMP-MAVGs encoded neither. This suggests that: (1) glutamine riboswitches are a common but not defining feature in Melnitz-like marine phages; (2) phages of that group use riboswitches or tmRNA, but rarely both, with the only two identified examples occurring in isolated phages, not metagenomically derived genomes.

Curiously, the bacterial OM43 host of Melnitz (H5P1) does not encode a glutamine riboswitch - only one cobalamin riboswitch was found located upstream of the Vitamin B12 transporter gene *btuB*. Other members of the *Methylophilaceae* (OM43 strains HTCC2181, KB13, MBRSH7 and *Methylopumilus planktonicus, M. rimovensis* and *M. turicensis*) also lack glutamine riboswitches. In cyanobacteria, the glutamine riboswitches were previously found to regulate the glutamine synthase *glnA* and are strongly associated with nitrogen limitation (61). Phages infecting *Synechococcus* have been shown to use extracellular nitrogen for phage protein synthesis (62), which might indicate a similar role for phage-encoded glutamine riboswitches in Melnitz. However, neither OM43 nor Melnitz encode the glutamine synthase *glnA* or homologous genes required for glutamine synthesis. This suggests that Melnitz-like phages use viral riboswitches to regulate their own genes rather than hijacking the cellular machinery.

### Phage encoded curli operons may be involved in regulating cell lysis

Curli genes are typically associated with the bacterial production of amyloid fibres that are part of biofilm formation, where the CsgGF pore spans the outer membrane as part of a Type VIII secretion system, allowing for the secretion of CsgA and CsgB that assemble into extracellular amyloid fibres (63). In complete curli modules, CsgG and CsgF form an 18-mer heterodimer comprising nine subunits with 1:1 stoichiometry between CsgF and CsgG. The structure of the pore is dictated by CsgG, which forms a channel ∼12.9Å in diameter. CsgF forms a secondary channel ∼14.8Å in diameter at the neck of the beta barrel (Figure 4A, Supplementary Figure 6 A and B) and assists in excretion of the amyloid fibre. Like previously described pelagimyophages (28), Melnitz possesses *csgF* and *csgG*, but lacks the genes for amyloid fibre production - *csgA* and *csgB* (Figure 3). Zaragoza-Solas et al. speculated that phage-encoded curli pores may allow for the uptake of macromolecules, or together with unidentified homologues of missing curli genes form a complete, functional curli operon producing amyloid fibres for “sibling capture” of proximate host cells (28). However, similar to results in pelagimyophages, we did not find evidence for a complete phage encoded curli biogenesis pathway in Melnitz, nor does the bacterial H5P1 host encode for any curli associated genes, suggesting that these genes were acquired by an ancestral strain of Melnitz and pelagimyophages before host transition to OM43 and SAR11, respectively.

**Figure 4.**
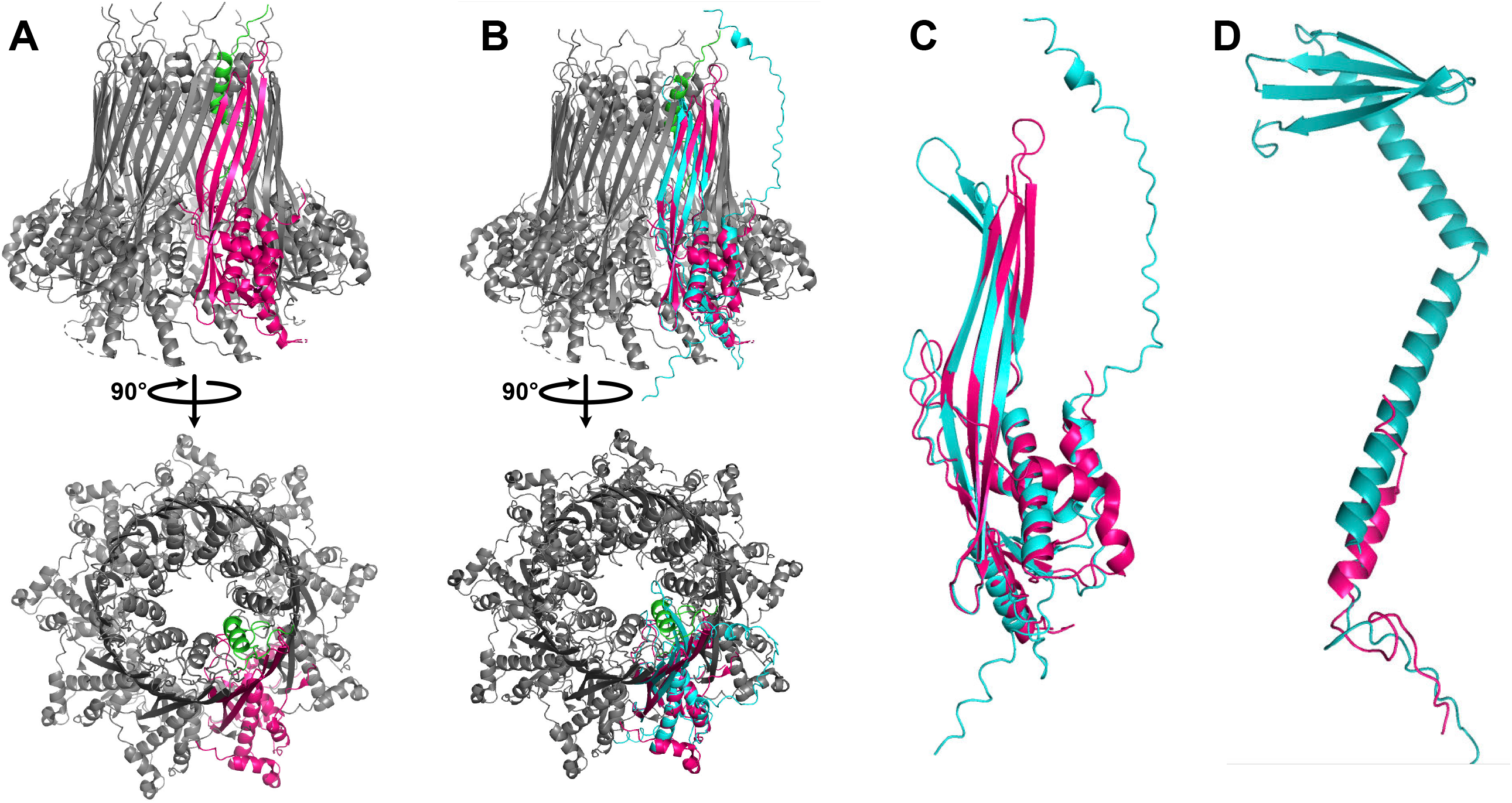
Structural prediction of CsgGF complex encoded by Melnitz. **A:** Predicted structure of CsgGF complex in *E. coli* (PDB model 6l7a) comprises a hetero-18-mer with 1:1 stoichiometry of CsgG (pink) and CsgF (green), forming a pore in the outer membrane with two constrictions - one provided by CsgG at the base of the barrel and one provided by CsgF at the neck of the barrel. **B:** Structural prediction of Melnitz encoded CsgG using AlphaFold2 (teal) showed structural conservation with CsgG from *E. coli* in the periplasmic α-helices and β-barrel structure. **C:** Expanded view of the structural alignment of Melnitz CsgG with *E. coli* CsgG shows a putative narrowing of the pore at the top of the barrel, matching the pore diameter at the top of the barrel in CsgGF in E. coli. **D:** Alignment of predicted structure of Melnitz-encoded CsgF (teal) to that of CsgF in *E. coli* (pink) showed low structural similarity, with Melnitz-encoded CsgF comprising two alpha-helices and a beta sheet. Alignment indicated that the additional structures of Melnitz-encoded CsgF extend out of the CsgG pore, with unknown function.

**Figure 5.**
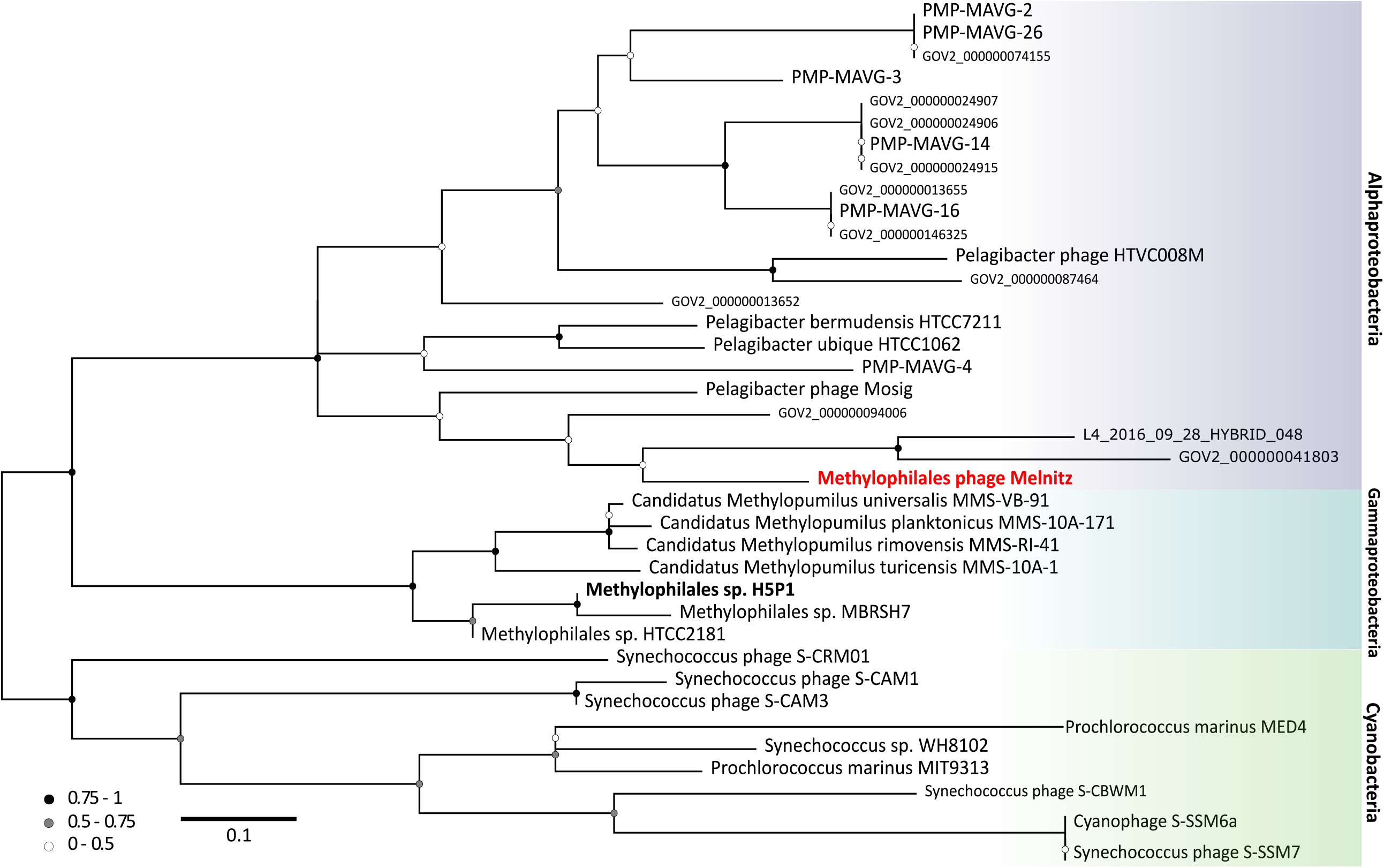
Phylogeny of tmRNA genes in major marine lineages. Neighbour-joining tree (100 bootstraps) of tmRNA genes found in marine phages and host lineages (not exhaustive) suggests that three the three known major lineages between Cyanobacteria, Gammaproteobacteria and Alphaproteobacteria are shared with their associated phages, except for OM43 phage Melnitz (infecting H5P1 on the Gammaproteobacterial branch) which has a tmRNA gene more closely related to genes found in Alphaproteobacteria and their phages.

Structural homology modelling of the Melnitz-encoded CsgGF portal with SwissModel (64) yielded low structural similarity to known CsgGF structures (GMQE <0.5 with Q-MEAN scores < -3 and z-scores >>2 outside normalised Q-mean score distributions from a non-redundant set of structures from PDB). Structural similarity was greatest in regions 238-258 of CsgG where Q-MEAN scores were ∼0.7, with conserved regions in the periplasmic-facing α-helices of CsgG. Melnitz-encoded CsgF showed poor structural similarity to any models available within Swiss-Model. Therefore, we used AlphaFold2 (65) to predict the structure of phage-encoded CsgG and CsgF *de novo* and compared predicted structures to known CsgGF structures. The predicted structure of Melnitz-encoded CsgG showed close structural similarity to known CsgG, with two notable exceptions: (1) a narrowing of the barrel at the neck; (2) an extension of the α-helix outside the barrel (Figure 4B and 4C). In contrast there was little structural similarity between Melnitz-encoded CsgF and the structure of CsgF in *E. coli*, barring a shared α-helix domain. Phage-encoded CsgF comprised two long α-helices ending in a β-sheet (Figure 4D). Similar modelling of CsgG and CsgF encoded by pelagiphage HTVC008M also contained these features (Supplementary Figure 7 A and B, respectively). Assuming the pore orientation in an infected OM43 cell is the same as that of CsgG in *E. coli*, the additional domains on phage-encoded CsgF would either place the terminal β-sheet inside the barrel of CsgG (unlikely due to steric hindrance) or the extended structure would point outwards into the extracellular milieu, in an unusual conformation with unknown function. It is more likely that in this configuration of CsgG, phage-encoded CsgG and CsgF have diverged and evolved independently. Therefore they may no longer form a heterodimer, with CsgG retaining the function as a pore, and CsgF evolving independently to provide an alternative, unknown function. Indeed, the narrowing of the barrel neck in Melnitz-encoded CsgG creates a channel 14.7Å in diameter, matching that provided by CsgF in *E. coli* (Supplementary figure 6 B and C). One possibility is that phage-encoded CsgG is functionally analogous to pinholins, used by phage λ to regulate cell lysis. Pinholins form channels ∼15Å in diameter in the inner membrane, resulting in rapid membrane depolarization and subsequent activation of membrane-bound lysins (66).

An alternative hypothesis is that in Melnitz, CsgGF forms a heterodimer in the outer membrane similar to CsgGF in *E. coli*, but that the structure is inverted so that the extended α-helices of CsgF point into the periplasm. In this conformation, the extension from Melnitz-encoded CsgG (Figure 4C) can interact to stabilise the elongated hinged α-helices of CsgF. The C-terminal β-sheets of CsgF could then form a second channel beneath the CsgG barrel (Supplementary Figure 8 A). In this conformation, CsgGF is structurally similar to a secretin, a large protein superfamily used for macromolecule transport across the outer membrane such as DNA for natural competence (e.g. PilQ within the type IV secretion system in *Vibrio cholerae*) or extrusion of filamentous phages during chronic infection (e.g. pIV in bacteriophages Ff) (67). Like CsgGF in an inverted conformation, PilQ comprises a β-barrel, with a secondary pore below, attached by two hinged α-helices. The inner surface of PilQ is negatively charged to repel the negatively charged backbone of DNA and assist in transportation across the membrane. In contrast, the inner surface of the CsgGF channel is positively charged and narrower (14.7Å compared to 20.6Å in PilQ), making a role in DNA transport across the membrane unlikely. While the function of phage-encoded CsgGF is not yet clear, the additional domains of CsgF and the lack of other curli genes within either the phage or the host suggest that it is a novel pore structure whose function has diverged from ancestral CsgGF and would be a worthy target for future structural resolution.

Whether the CsgGF complex in Melnitz acts as a pinholin or a secretin, gene synteny supports a putative role in regulation of the timing of cell lysis. First, genes *csgGF* are located immediately downstream of thymidylate synthase *thyX* and *ssrA* and transcription co-activator genes (Figure 3). Overexpression of thymidylate synthase in the D29 phage infecting *Mycobacteria tuberculosis* results in delayed lysis and higher phage yields (68). In phage T4, postponing lysis is used to delay the release of viral progeny until conditions are favourable, thereby maximising virion production before viral release and successful replication after (69). In the T4-like Melnitz, *thyX* is therefore likely to be involved in postponing lysis as well. Lytic control may putatively be related to the upstream glutamine riboswitch. H5P1 lacks glutamine synthesis pathways, but has a complete peptidoglycan biosynthesis pathway necessary to produce peptidoglycans from glutamine. Therefore, both the H5P1 host and Melnitz phage are restricted to using glutamine for protein and peptidoglycan synthesis only. In phage λ, the depletion of peptidoglycan precursors can trigger lysis through activation of a spanin complex (70). We speculate that during the course of the infection cycle, glutamine levels are kept low through incorporation into viral proteins. When protein synthesis is complete, intracellular glutamine levels increase, activating the glutamine riboswitch. This in turn could trigger an opening of the curli portal, which may result in rapid depolarization of the membrane, triggering activation of lysins and rapid cell lysis.

Structural homology modelling (SwissModel) of a Melnitz-encoded enzyme (gp67), putatively annotated as a glycosyl hydrolase, revealed structural similarity (38% sequence identity at 98% coverage, with a global model quality estimate (GMQE) of 0.82) to autolysin SagA encoded by *Brucella abortus* (PDB model 7DNP). SagA acts to generate localised gaps in the peptidoglycan layer for assembly of type IV secretion systems (71). Therefore, it is likely that gp67 and CsgGF work in concert in Melnitz. Peptides involved in binding peptidoglycan at the active site (Glu17, Asp26, Thr31) were conserved between Melnitz gp67 and SagA (Supplementary Figure 9X) (72). Outside of this active site, structural similarity to T4 endolysin (PDB model 256l.1) was low (13% identity over 51% coverage; GMQE: 0.18). Melnitz lacked any other lysin-like genes. Like the T4-encoded endolysin *e* (73), gp67 is under control of an early transcription promotor (Figure 3B). We therefore propose that the gp67 gene in Melnitz potentially serves two functions: (1) as an endolysin for degradation of cellular peptidoglycans during lysis; or (2) as an autolysin to enable assembly of the CsgGF or CsgG-only portal protein. Phages infecting *Streptococcus pneumoniae* upregulate host-encoded autolysins alongside phage-encoded lysins during late-stage infection to accelerate cell lysis (74). The close structural match between gp67 and SagA suggests that an ancestor of Melnitz acquired a host-encoded autolysin as an alternative lysin during its evolutionary history. Whether this autolysin-derived phage lysin could be activated by membrane depolarization through CsgG is unknown.

### Melnitz-encoded genes suggest a possible host transition event from SAR11 to OM43

Phage host ranges are largely determined by interactions between host receptor proteins and phage structural proteins such as tail fibres that enable the phage to adsorb and inject its genetic material. Mutation in phage proteins, either through point mutations or recombination during co-infection can result in host range expansion or transition (75). Host range expansion or transition within species boundaries is more common, but rare transition events between hosts from different genera can occur through mutations in tail fibres (76). Here, two separate lines of evidence converge to suggest that Melnitz underwent a recent host transition from SAR11 to OM43. First, *ssrA* encoded by Melnitz was identified to be more closely related to homologs within the alphaproteobacterial lineage. Two-piece circularly permuted tmRNA is common in three major lineages: Alphaproteobacteria, Betaproteobacteria (now part of Gammaproteobacteria (77)) and Cyanobacteria (78, 79). The phylogenetic evidence for alphaproteobacterial *ssrA* in Melnitz rather than *ssrA* that matches the gammaproteobacterial lineage of its OM43 host suggests that this gene was acquired from an Alphaproteobacteria (Figure 4). Second, two of the four tRNAs encoded by Melnitz were more closely related to tRNAs encoded by pelagimyophages than those of their host. Melnitz encodes two versions of tRNA-Arg(TCT) – one most closely related to that of its host H5P1, and one closely related to *Pelagibacter* phage Mosig. Phage-encoded tRNAs enable large phages to sustain translation as the host machinery is degraded to fuel phage synthesis (80). Phage-encoded tRNAs also enable phages to optimise protein synthesis in a host with different codon usage and thus serve as both a marker of increased host range and evolution through different hosts (81). Indeed, tRNAs are often used for computational host-prediction of phages due to high sequence conservation of the gene between host and viral forms (82, 83). We postulated that the host-range of Melnitz would be evident in the four tRNAs found within its genome and reflect potential hosts in the OM43 clade. Alternatively, if a recent host transition occurred, as *ssrA* phylogeny suggests, tRNAs would be similar to those found in the SAR11 bacteria and viruses. A search for tRNA genes in the genomes of isolated *Pelagibacter* phages and PMP-MAVGs (47 genomes) and isolated phages infecting OM43 (three genomes) identified tRNAs in seven *Pelagibacter* phage genomes. Melnitz was the only phage known to infect OM43 that encoded tRNAs, with four tRNAs in total (Figure 6). Sequences encoding tRNAs were aligned using all-vs-all BLASTN with a minimum expect-value of 1 × 10^−5^. Two out of four tRNAs in Melnitz were tRNA-Arg(TCT), the first aligning most closely with its H5P1 host. The second tRNA-Arg best aligned with the tRNA gene found in the *Pelagibacter* phage Mosig. Its alignment to bacterial tRNA matched OM43 strains H5P1 and HTCC2181, respectively (>90% identity). The third tRNA-Leu found in Melnitz aligned with PMP-MAVG-17, previously classified as *Pelagibacter* phage (28), but only had 45% nucleotide identity. The fourth tRNA-Trp (CCA) found in Melnitz did not align (e-values > 1 × 10^−5^) with any tRNA found in OM43, SAR11 or any of their respective phages. Using tRNA as host indication in Melnitz therefore reflects its OM43 host, but also indicates genetic exchange with SAR11 virus-host systems. Furthermore, the related *Pelagibacter* phage Mosig possessed tRNAs matching OM43 and Melnitz as well as a second tRNA aligning with its SAR11 host (93%). This may suggest either a relatively recent genetic exchange between OM43 and SAR11 virus-host systems, and/or a surprisingly broad host-range for *Pelagibacter* phage Mosig and *Methylophilales* phage Melnitz.

**Figure 6.**
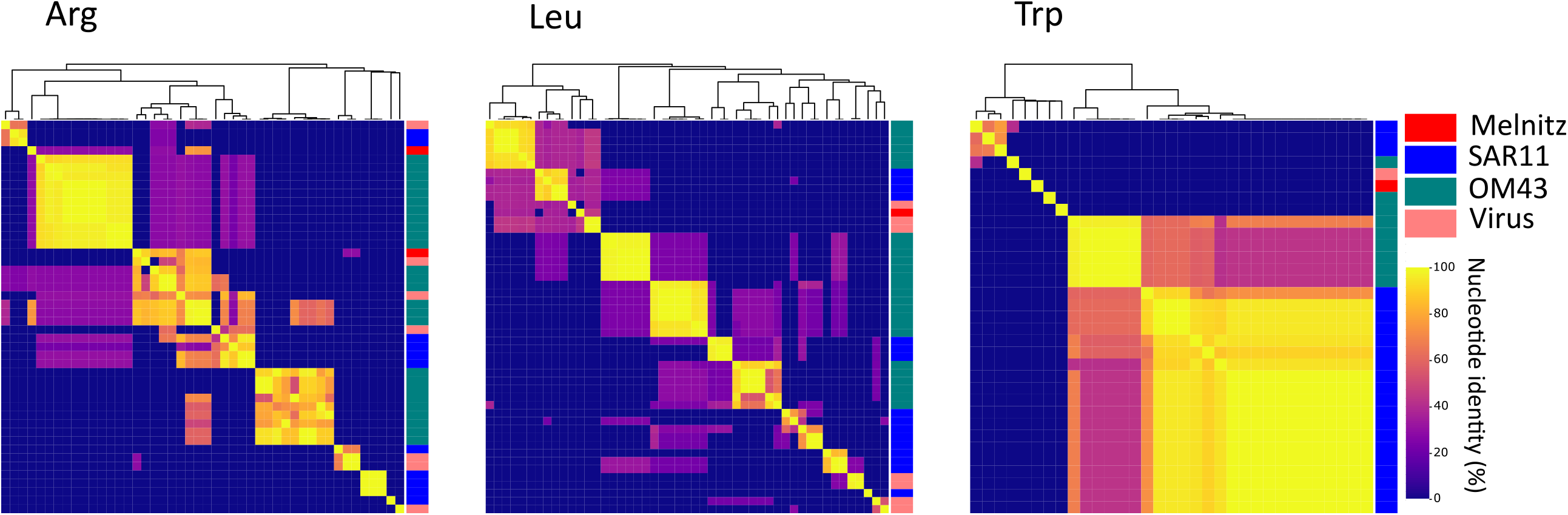
Alignment of tRNA genes found in Melnitz, SAR11 and OM43 lineages. Heatmaps and dendrograms (Euclidian similarity matrices) based on similarity between alignments for arginine (Arg), leucine (Leu) and tryptophan (Trp) tRNA genes found in OM43 phage Melnitz, OM43 and SAR11 as well as tRNA derived from isolated SAR11 and OM43 phages.

To assess the breadth of host ranges, we challenged the SAR11 strains HTCC7211 and HTCC1062 and OM43 strains H5P1, D12P1 and C6P1 (15) with the phages Melnitz and Mosig (27), but found no evidence that either phage is able to replicate in cells beyond host class boundaries as observed via cell lysis (Supplementary Figure 10). Melnitz only infected D12P1 and H5P1, but not C6P1 or any SAR11, whereas Mosig only caused lysis in the *Pelagibacter ubique* HTCC1062. Though the limited number of available bacterial strains could have missed potential permissive hosts in a different subclade, there is no evidence that non-canonical matching tRNA between SAR11/OM43 and their phages is due to a speculative broad host range in these systems, and instead supports the phylogenetic evidence we presented for a recent host transition event.

Phage host-range expansion and transition within closely related (i.e. between strains) is a common feature of phage evolution, and has been shown to occur *in vitro* via homologous recombination between co-infecting phages (75, 84). In contrast, expansion and transition between distantly related taxa is rare but has been previously observed *in vitro*. Naturally occurring, low-abundance mutants of T4 transitioned from *E. coli* to *Yersinia pseudotuberculosis* through modification of tail fibre tips in gene 37, yielding variants that could infect both hosts and others that could only infect *Y. pseudotuberculosis* (76). To our knowledge, host transition between bacteria belonging to different classes, as proposed here, has not previously been observed, although host-prediction of viral contigs from metagenomes has identified rare phages (115 out of 3,687) possibly able to infect multiple classes (85). We propose that two properties of SAR11 and OM43 increase the likelihood of such an event in natural communities: (1) Both SAR11 hosts and their associated phages possess extraordinarily large, globally ubiquitous effective population sizes, making even extreme rare events likely. This study has shown that phages infecting OM43 can also be abundant at higher latitudes, further increasing the likelihood of rare strain variants to occur; (2) Both OM43 and SAR11 hosts share highly streamlined genomes with elevated levels of auxotrophy, minimum regulation and similar G+C content, shaped by selection pressure to maximise replication on minimum resources in nutrient-limited marine environments (19, 86). Both fastidious hosts can be cultured on identical minimal medium, as long as additional methanol is provided as a carbon source for the methylotrophic OM43 (15). Therefore, the phenotypic difference between OM43 and SAR11 might be smaller than suggested by their taxonomic classification. Given viral selection occurs at the phenotypic level, we propose the possibility of a rare event where a mutant of a T4-like phage infecting SAR11 was able to successfully adsorb and inject its genome into an OM43 host, possibly co-infected with a methylophage that enabled a host-transition event through homologous recombination. Once a transition event occurred, the phage likely rapidly evolved specialism on the new host, losing the ability to infect the original host, as demonstrated in host-range experiments. Compared to copiotrophic, *R*-strategist microbial taxa such as *E. coli* and *Vibrio* spp., very little is known about the processes governing viral host-range and co-evolution in *K*-strategist bacteria such as the genomically streamlined taxa that dominate oligotrophic oceans. The early evidence shown here may suggest that phages can transition to new hosts from distantly related taxa in natural communities. Such events would explain in part the diverse host ranges predicted amongst viruses within gene-sharing clusters (87, 88).

## Conclusion

In this study, we provide evidence that supports a putative inter-class host transition event between two important clades of streamlined marine heterotrophs and expand our knowledge about the dynamics and characteristics in genomically streamlined heterotrophic virus-host systems. We isolated four near-identical strains of the new myophage Melnitz from subtropical and temperate marine provinces infecting the important methylotrophic OM43 clade, which we showed to be closely related to myophages infecting the abundant SAR11 clade. The analysis of metagenomic datasets provides evidence that this phage group is ubiquitous in global oceans despite relatively low overall abundance, supporting the viral seed bank hypothesis. Our genomic analysis of Melnitz revealed an incomplete curli module similar to reported curli pores in pelagimyophages, representing a rare and intriguing protein dimer that is absent in their respective host clades. We propose that these virally encoded curli pores may have been repurposed as a functional analogue for the regulation of viral lysis. We also identified an *ssrA* gene encoding for a complete viral tmRNA protein controlled by a glutamine riboswitch, showing that virus-host interactions can be regulated through riboswitches reflecting the extensive use of riboswitches in streamlined marine heterotrophs. Further phylogenetic analysis showed that the *ssrA* gene is related to the alphaproteobacterial SAR11 lineage, not the gammaproteobacterial OM43 lineage, providing evidence for host transition events in natural marine microbial communities, which was supported by the alignment of viral and bacterial tRNA genes of both lineages. These findings support the conclusion that in heterotrophic streamlined virus-host systems evolution of viral diversity is likely to be driven by host transition and expansion between closely related phages infecting hosts across broad taxonomic groups, likely increasing mosaicism and genetic exchange.

## MATERIALS & METHODS

### OM43 strain, media, and growth conditions

The OM43 strains *Methylophilales* H5P1 and D12P1 were isolated previously from surface water from the Western English Channel (15). Continuous cultures were grown using artificial seawater-based artificial seawater medium (ASM1) (89) amended with 1 mM NH_4_Cl, 10 µM KH_2_PO_4_, 1 µM FeCl_3_, 100 µM pyruvate, 25 µM glycine, 25 µM methionine as well as 1 nM each of 4-amino-5-hydroxymethyl-2-methylpyrimidine (HMP), pantothenate, biotin, pyrroloquinoline quinone (PQQ) and B12. Additional 1 mM methanol and 5 µL of an amino acid mix (MEM Amino Acids (50x) solution, Sigma-Aldrich) were added per 100 mL medium. Bacterial stocks were grown in 50 mL acid-washed polycarbonate flasks at 15 °C without shaking and 12-hour light-dark cycles.

### Water sources and phage isolation

Water samples were collected from two different stations: Station L4 in the Western English Channel (WEC; 50°15’N; 4°13’W) and the Bermuda Atlantic Time Series sampling station (BATS; 31°50’N; 64°10’W) in the Sargasso Sea. For each sample we used Niskin bottles mounted on a CTD rosette to collect 2 L of seawater at 5m depth (Table 1) into acid-washed, sterile, polycarbonate bottles. To obtain a cell-free fraction, water samples were filtered sequentially through a 142 mm Whatman GF/D filter (2.7 µm pore size), a 142 mm 0.2 µm pore polycarbonate filter (Merck Millipore) and a 142 mm 0.1 µm pore polycarbonate filter (Merck Millipore) using a peristaltic pump. The cell-free viral fraction was concentrated to about 50 mL with tangential flow filtration (50R VivaFlow, Sartorius Lab instruments, Goettingen, Germany) and used as inoculum in a multi-step viral enrichment experiment followed by Dilution-to-Extinction purification as described previously (15). Briefly, viral inoculum (10% v/v) was added to 96-well Teflon plates (Radleys, UK) with exponentially growing host cultures. After a 1 to 2 weeks incubation period, cells and cellular debris were removed with 0.1 µm syringe PVDF filters. The filtrate was added as viral inoculum to another 96-well Teflon plate with exponentially growing host culture. This process was repeated until viral infection was detected by observing cell death using flow cytometry. Phages were purified by Dilution-to-Extinction methods (Our detailed protocol is available here: 10.17504/protocols.io.c36yrd).

### Assessment of viral host ranges

An acid-washed (10% hydrochloric acid) 48-well Teflon plate was prepared with 2 mL per well of ASM1 amended with 1 mM methanol and 5 µL of an amino acid mix (MEM Amino Acids (50x) solution, Sigma-Aldrich). Wells were inoculated to 1×10^6^ cells·mL^-1^ with *Pelagibacter* hosts HTCC1062 or HTCC7211, or OM43 hosts C6P1, D12P1 and H5P1 cultures in replicates of three for each of the Mosig and Melnitz phages, plus one set of no virus controls for each host. The wells marked for viral infections were infected with 200 µL of viral culture. The plate was then incubated at 15 °C and monitored daily using flow cytometry for ∼2 weeks. Successful infections were identified observing cell lysis in virus amended wells compared to no-virus controls (15).

### Phage DNA preparation, genome sequencing and annotation

For each viral isolate, 50 mL OM43 host cultures were grown in 250 mL acid-washed, polycarbonate flasks and infected at a cell density of about 1-5x 10^6^ cells·mL^-1^ with 10% v/v viral inoculum. Infected cultures were incubated at 15 °C for 7 to 14 days after which the lysate was transferred to 50 mL Falcon tubes. Larger cellular debris was removed by centrifugation (GSA rotor, Thermo Scientific 75007588) at 8,500 rpm/ 10,015 x g for one hour. Supernatant was filtered through pore-size 0.1 µm PVDF syringe filter membranes to remove any remaining smaller cellular debris. Phage particles were precipitated using a PEG8000/NaCl flocculation approach (90). Briefly, 50 mL lysate was amended with 5g PEG8000 and 3.3 g NaCl (Sigma), shaken until dissolved and incubated on ice overnight. Precipitated phages were pelleted by centrifugation at 8,500 rpm / 10,015 x g for one hour at 4 °C. After discarding the supernatant, phage particles were resuspended by rinsing the bottom of the tubes twice with 1 mL SM buffer (100 mM NaCl, 8 mM MgSO_4_·7H_2_O, 50 mM Tris-Cl). DNA from the suspended phages was extracted with the Wizard DNA Clean-Up system (Promega) following manufacturer’s instructions, using pre-heated 60 °C PCR-grade nuclease free water for elution.

Nextera XT DNA libraries were prepared and sequenced by the Exeter Sequencing Service (Illumina paired end [2 × 250 bp], NovaSeq S Prime [SP], targeting 30-fold coverage). Reads were quality controlled, trimmed and error corrected with the tadpole function (default settings) within BBMap v38.22 ((91) available at https://sourceforge.net/projects/bbmap/). Contigs were assembled using SPAdes v.3.13 with default settings (92). Viral contigs were confirmed with VirSorter v1.05 (93) (categories 1 or 2, > 15kbp). Quality and completeness of contigs as well as terminal repeats were evaluated using CheckV v0.4.0 with standard settings (94). Genes were called with phanotate v2019.08.09 (95) and imported into DNA Master for manual curation (96) using additional gene calls made by GenMarkS2 (97), GenMark.heuristic (98), Prodigal v2.6.3 (99) and Prokka v1.14.6 (100). ORFs were functionally annotated using BLASTp against NCBI’s non-redundant protein sequences (101), phmmer v2.41.1 against Pfam (102) and SWISS PROT (103). All genes called were listed and compared using a scoring system evaluating length and overlap of ORFs as well as quality of annotation (96); tRNA and tmRNA were identified with tRNAScan-SE v2.0 (104) and ARAGORN v1.2.38 (59). Genomes were scanned for riboswitches using the web application of Riboswitch Scanner (106, 107). FindTerm (energy score < -11) and BPROM (LDF > 2.75) from the Fgenesb_annotator pipeline (108) were used to predict promoter and terminator sequences, using default parameters. The σ^70^ promoters predicted this way were considered early promoters. Known T4-like late promoter sequence 5’-TATAAAT-3’ (50, 56) and middle-promoter MotA box (TGCTTtA) dependent middle promoters were used as query for a BLASTN search over the whole genome. Promoters and/or terminators were excluded if they were not intergenic or not within 10 bp overlap of the start/end of ORFs.

### Host DNA preparation, genome sequencing and annotation

Host cultures were grown in 50 mL ASM1 medium amended with 1mM methanol in 250 mL polycarbonate flasks. Upon reaching maximum cell density, genomic DNA was extracted using Qiagen DNAeasy PowerWater Kit (REF 14900-50-NF) from biomass retained on 0.1 µm PC filters following the manufacturer’s protocol with minor modifications to increase the yield. The bead beating step was lengthened from 5 minutes to 10 minutes. DNA elution was performed with a 2 minute incubation with elution buffer warmed to 55 °C. Nextera DNA libraries were prepared and sequenced by MicrobesNG (Birmingham, UK) for Illumina short read sequencing on the HiSeq2500 (Illumina paired end [2 × 250 bp], targeting 30-fold coverage). Additionally, long read sequencing was prepared using a MinION flow cell. Reads were quality controlled, trimmed and error corrected with the tadpole function (default settings) within BBMap v38.22 ((91) available at https://sourceforge.net/projects/bbmap/). Contigs were assembled using SPAdes v.3.13 with default settings (92). Gene calls were made with PROKKA v1.14.6 (100) and submitted to BlastKOALA (105) for further annotation and prediction of KEGG pathways using the *Methylophilales* strain HTCC2181 as reference.

### Phylogeny and Network analysis

All contigs from the Global Ocean Virome (GOV2) dataset and a WEC virome (12, 29), *Methylophilales* phage Venkman (15) and LD28 phage P19250A (109), all isolated *Pelagibacter* phages (15–18, 27) and *Pelagibacter*-like MAGs (28) were screened for contigs that share a viral population with the genomes of OM43 phages (95% ANI over 80% length) using ClusterGenomes.pl v5.1 (https://github.com/simroux/ClusterGenomes). Genes of all contigs were identified by Prodigal v2.6.3 (99) and imported into the Cyverse Discovery Environment 2.0 (available at https://de.cyverse.orge/), where vContact-Gene2Genome 1.1.0 was used to prepare protein sequences before protein clustering using VConTACT2 (v.0.9.8) with default settings (87) to assess relatedness via shared gene networks. The gene sharing network was visualised using Cytoscape v3.7.1 (110). For the single-gene based phylogeny, genes of contigs that fell into the same viral protein clusters as the isolated OM43 phages from this study were aligned to selected genes in annotated OM43 phage genomes, and other respective genes of interest, with BLASTp (default parameters) (111). Genes were aligned within the Phylogeny.fr online server (112) opting for MUSCLE alignment (113) and built-in curation function (114) with default settings, removing positions with gaps for calculating phylogenetic trees. Maximum-likelihood trees were calculated with PhyML (115, 116) using 100 bootstraps. The trees were visualised using FigTree (v1.4.4 available at http://tree.bio.ed.ac.uk/software/figtree/). Figures were edited with Inkscape (www.inkscape.org) for aesthetics.

### Metagenomic reads recruitment

Marine virome datasets were used to assess the relative abundance of phage contigs including a single virome from the Western English Channel and 131 samples from the Global Ocean Virome dataset (GOV2) (12, 29). Metagenomic reads were subsampled to 5 million reads using the reformat.sh command within the bbmap suite. Bowtie2 (117) indexes of dereplicated contigs were created for all known pelagiphage isolate genome (15–18, 27), the LD28 phage P19250A (109), a selection of cyanophages, a selection of abundant *Roseobacter* phages (118), *Enterobacteria* phages T4 and T7 (as negative controls) as well as one genome from the viral population isolated in this study (Melnitz). Metagenomic reads from each virome were mapped against all contigs with bowtie2 (bowtie2 --seed 42 --non-deterministic). To calculate coverage and Reads Per Kilobase of contig per Million reads (RPKM) we used coverm (available at https://github.com/wwood/CoverM), with the following commands: “coverm contig --bam-files *.bam --min-read-percent-identity 0.9 --methods rpkm --min-covered-fraction 0.4”.

### Search and alignment of tRNA

tRNAs were identified from bacteria and virus genomes using ARAGORN v.1.2.38 (59) using the ‘-t – gcbact –c –d –fons’ flags and tRNAscan-SE v2.0.7 using flags ‘-B –fasta’ (104). tRNAs were deduplicated using seqkit v0.15.0 (119) with ‘rmdup –by-seq –ignore-case’. A BLAST database of tRNA genes was made in BLAST v2.5.0+ (111) and used for sequence alignment with BLASTN with the flags ‘-outfmt ‘6 std qlen slen’ -evalue 1e-05 -task blastn-short’. Percentage identity was calculated by dividing alignment-length by query-length times alignment-percentage.

### Structural analysis of a putative endolysin

The predicted amino acid sequence of gene product 67 (gp67) was used as a query to identify putative structures on the SwissModel server (64) using BLAST and HHBits. Putative models were downselected based on suitable quaternary structure properties and GQME>0.7. Autolysin SagA from *Brucella abortus* (PDB model 7dnp.1) was selected as the best-hit and used for subsequent modelling of the structure of gp67. Models and associated figures were visualised in PyMOL v. 2.5.1 (https://pymol.org/2/).

### Structural analysis of CsgGF

The predicted amino acid sequences of CsgG and CsgF from Melnitz were used as a query to identify putative structures on the SwissModel server using BLAST and HHBits. The best-hit was determined by predicted quaternary structure properties. Global GQME scores were <0.5 and Q-MEAN scores identified the best predicted structures as unreliable (ranging from -3.28 to -5.11), although localised regions had Q-MEAN scores > 0.7. Therefore, to improve structural predictions, amino acid sequences for CsgG and CsgF were independently run through AlphaFold2 using the available Colab web interface (https://colab.research.google.com/drive/1LVPSOf4L502F21RWBmYJJYYLDlOU2NTL). Structures were determined both with and without post-prediction relaxation (use_amber) and use of MMSeqs2 templates (use_templates). As no noticeable differences were observed between these runs, we selected the top scoring unrelaxed model for downstream comparison to known CsgG and CsgF structures. Predicted structures from AlphaFold2 were downloaded, visualised and aligned to *E. coli* CsgGF (7NDP) in PyMOL v. 2.5.1. CsgGF from pelagimyophage HTVC008M was similarly analysed with AlphaFold2 to confirm structural similarity to that of Melnitz. Structural prediction of the CsgGF heterodimer in Melnitz was assumed to conform to the 18-mer structure of CsgGF in *E. coli*. Scripts for generating structures in PyMOL are available here https://github.com/HBuchholz/Genomic-evidence-for-inter-class-host-transition-between-streamlined-heterotrophs. Electrostatic potential of protein surfaces was calculated and visualised using the APBS Electrostatics plugin available within PyMOL.

### Transmission Electron Microscopy

For ultrastructural analysis, bacterial cells and/or phages were adhered onto pioloform-coated 100 mesh copper EM grids (Agar Scientific, Standsted, UK) by floating grids on sample droplets placed on parafilm for 3 min. After 3 × 5 min washes in droplets of deionized water, structures were contrasted on droplets of 2% (w/v) uranyl acetate in 2% (w/v) methyl cellulose (ratio 1:9) on ice for 10 min, the grids picked up in a wire loop and excess contrasting medium removed using filter paper. The grids were then air dried, removed from the wire loop and imaged using a JEOL JEM 1400 transmission electron microscope operated at 120 kV with a digital camera (Gatan, ES1000W, Abingdon, UK).

## DATA AVAILABILITY

All four Melnitz-like genome were deposited as GenBank entry on NCBI under accession number MZ577095-MZ577098 of BioProject PRJNA625644; the reference genome used for the analysis was deposited under MZ577097. Sequencing data for all phages sequenced in this study can be found on the SRA databank under Accession numbers SAMN18926670-SAMN18926674. Reads for Methylophilales bacterial host H5P1 are available under SAMN20856461.

## ACKNOWLEDGEMENTS

We would like to thank Christian Hacker and the Bioimaging Centre of the University of Exeter for performing the TE microscopy and imaging. We would also like to thank the crew of the R/V Plymouth Quest and our collaborators at Plymouth Marine Laboratory for collecting water samples, and the driver Magic for delivering water samples from Plymouth to Exeter. We would also like to thank the crew of the R/V Atlantic Explorer and our collaborators at the Bermuda Institute of Ocean Sciences. We acknowledge the use of the University of Exeter High-Performance Computing (HPC) facility in carrying out this work. Genome sequencing was provided by the Exeter Sequencing Service. This project utilised equipment funded by the Wellcome Trust Institutional Strategic Support Fund (WT097835MF), Wellcome Trust Multi User Equipment Award (WT101650MA) and BBSRC LOLA award (BB/K003240/1). The efforts of Holger H. Buchholz in this work were funded by the Natural Environment Research Council (NERC) GW4+ Doctoral Training program. LMB, MLM and BT were funded by NERC (NE/ R010935/1) and by the Simons Foundation BIOS-SCOPE program.

H.H.B. and B.T. designed experiments and wrote the manuscript, H.H.B. performed experimental research and analysed data, B.T. performed protein structure predictions, L.M.B. assisted with data analysis, A.G.B. assisted with tRNA analysis, M.L.M. assisted with laboratory work.

---

Supplementary Figure 1. TEM images of phage Melnitz virions. Bar indicates 100 nm.

Supplementary Figure 2. Neighbour-joining tree (100 bootstraps) of a tail sheath encoding gene found in Melnitz related myophages. Branch support values <1 are indicated by circle colours on the tree. The cyanophage branch was used to root the tree.

Supplementary Figure 3. Neighbour-joining tree (100 bootstraps) of the *TerL* gene found in Melnitz related myophages. Branch support values <1 are indicated by circle colours on the tree. The cyanophage branch was used to root the tree.

Supplementary Figure 4. Neighbour-joining tree (100 bootstraps) of a capsid scaffolding gene found in Melnitz related myophages. Branch support values <1 are indicated by circle colours on the tree. The cyanophage branch was used to root the tree.

Supplementary Figure 5. Global abundance of Melnitz. Reads recruited per kilobase of contigs per million reads (RPKM) of GOV2 viromes against phages infecting OM43 and LD28 as well as known *Pelagibacter* myophages. Samples are organized by ecological zone: Arctic, TT-EPI temperate-tropical epipelagic, TT-MES temperate-tropical mesopelagic, ANT Antarctic.

Supplementary Figure 6. Cross sectional structure of CsgGF. **A** Clipped predicted surface model of internal structure of CsgGF in *E. coli* showing CsgG (teal) and CsgF (pink) showing the internal channel structure. This structure comprises a series of narrowing pores. **B** with CsgF creating a pore 14.8Å in diameter and CsgG creating a smallest pore 12.9Å in diameter. **C** Clipped predicted surface model of internal structure of Melnitz-encoded CsgG comprises two channels of similar size to those of the *E. coli* CsgGF complex.

Supplementary Figure 7. Structural comparison between **A** CsgG encoded by Melnitz (teal) and HTVC008M (pink); **B** CsgF encoded by Melnitz (teal) and HTVC008M (pink).

Supplementary Figure 8. **A** Predicted structure of Melnitz CsgF in an inverted orientation compared to **B** structure of secretin PilQ (6W6M) from *Vibrio cholerae*. Left to right - cartoon model; electrostatic potential (red = -ve, blue=+ve); top-down view through the barrel.

Supplementary Figure 9. SwissModel alignment of putative endolysin structure encoded by Melnitz gp67 (hot pink), with a SagA autolysin encoded by Brucella abortus (PDB model 7dnp.1, grey). The conserved peptidoglycan binding site encoded by Glu17, Asp26 and Thr31 is highlighted (teal), with the peptidoglycan substrate (yellow stick representation).

Supplementary Figure 10. Host range assessment of Melnitz. **A** Growth curve of SAR11 strains HTCC1062 and HTCC7211, treated with OM43 phage Melnitz and SAR11 phage Mosig. **B** Growth curves of OM43 strains C6P1, D12P1 and H5P1 treated with OM43 phage Melnitz and SAR11 phage Mosig

